# Detecting clinically actionable variants in the 3’ exons of *PMS2* via a reflex workflow based on equivalent hybrid capture of the gene and its pseudogene

**DOI:** 10.1101/379693

**Authors:** Genevieve M. Gould, Peter V. Grauman, Mark R. Theilmann, Lindsay Spurka, Irving E. Wang, Laura M. Melroy, Robert G. Chin, Dustin H. Hite, Clement S. Chu, Jared R. Maguire, Gregory J. Hogan, Dale Muzzey

## Abstract

**Background:** Hereditary cancer screening (HCS) for germline variants in the 3’ exons of *PMS2*, a mismatch repair gene implicated in Lynch syndrome, is technically challenging due to homology with its pseudogene *PMS2CL*. Sequences of *PMS2* and *PMS2CL* are so similar that next-generation sequencing (NGS) of short fragments—common practice in multigene HCS panels—may identify the presence of a variant but fail to disambiguate whether its origin is the gene or the pseudogene. Molecular approaches utilizing longer DNA fragments, such as long-range PCR (LR-PCR), can definitively localize variants in *PMS2*, yet applying such testing to all samples can have logistical and economic drawbacks.

**Methods:** To address these drawbacks, we propose and characterize a reflex workflow for variant discovery in the 3’ exons of *PMS2*. We cataloged the natural variation in *PMS2* and *PMS2CL* in 707 samples and designed hybrid-capture probes to enrich the gene and pseudogene with equal efficiency. For *PMS2* exon 11, NGS reads were aligned, filtered using gene-specific variants, and subject to standard diploid variant calling. For *PMS2* exons 12-15, the NGS reads were permissively aligned to *PMS2*, and variant calling was performed with the expectation of observing four alleles (i.e., tetraploid calling). In this reflex workflow, short-read NGS identifies potentially reportable variants that are then subject to disambiguation via LR-PCR-based testing.

**Results:** Applying short-read NGS screening to 299 HCS samples and cell lines demonstrated >99% analytical sensitivity and >99% analytical specificity for single-nucleotide variants (SNVs) and short insertions and deletions (indels), as well as >96% analytical sensitivity and >99% analytical specificity for copy-number variants. Importantly, 92% of samples had resolved genotypes from short-read NGS alone, with the remaining 8% requiring LR-PCR reflex.

**Conclusion:** Our reflex workflow mitigates the challenges of screening in *PMS2* and serves as a guide for clinical laboratories performing multigene HCS. To facilitate future exploration and testing of *PMS2* variants, we share the raw and processed LR-PCR data from commercially available cell lines, as well as variant frequencies from a diverse patient cohort.

## Background

Individual genomic variants inherited through the germline account for approximately 5% to 10% percent of cancer [1–3]. This heritable component can increase risk for malignancies across a range of tissues [4,5]—such as breast, colorectal, pancreatic, and prostate—and is associated with pathogenic variants in >100 genes [6]. To assess patients’ risk for such cancers, hereditary cancer screening (HCS) typically uses targeted next-generation sequencing (NGS) to detect relevant variants in the coding regions and select noncoding regions on a multigene testing panel.

In most genomic regions interrogated by HCS panels, NGS alone is sufficient to yield high sensitivity and specificity [7,8]; high accuracy is critical for HCS because test results prompt patients to alter their clinical-management decisions [9,10]. In a minority of regions, however, standard NGS strategies that use hybridization to capture and sequence short DNA fragments could incorrectly identify genotypes. Genes that pose particular challenges often have homologous sequences (e.g., pseudogenes) elsewhere in the genome that are captured and sequenced along with the gene itself, complicating alignment and the identification of variants specific to the gene.

*PMS2* is commonly included on HCS panels due to its association with Lynch syndrome [11–15]. Its nearby pseudogene, *PMS2CL*, complicates accurate NGS read alignment and variant identification in exons 11 through 15 at the 3’ end of *PMS2* (Fig. 1A): the coding sequences were previously reported to share 98% sequence identity with *PMS2CL* [16]. Further, sequence exchange and gene conversion between the two regions are sufficiently frequent that even the few non-identical bases in the reference genome (hg19) cannot be reliably attributed to the gene or pseudogene [17,18]. Long-range PCR (LR-PCR) using a gene-specific primer in exon 10 amplifies *PMS2* specifically (Fig. 1B), and variants in the terminal five exons of *PMS2* can then be identified via Sanger sequencing [19–21] or NGS [22] (Fig. 1C). Although identification of copy-number variants (CNVs) in *PMS2* is possible from LR-PCR and Sanger sequencing, it is not straightforward, which has motivated parallel use of multiplex ligation-dependent probe amplification (MLPA) to detect large deletions and duplications [19–24].

**Fig. 1.**
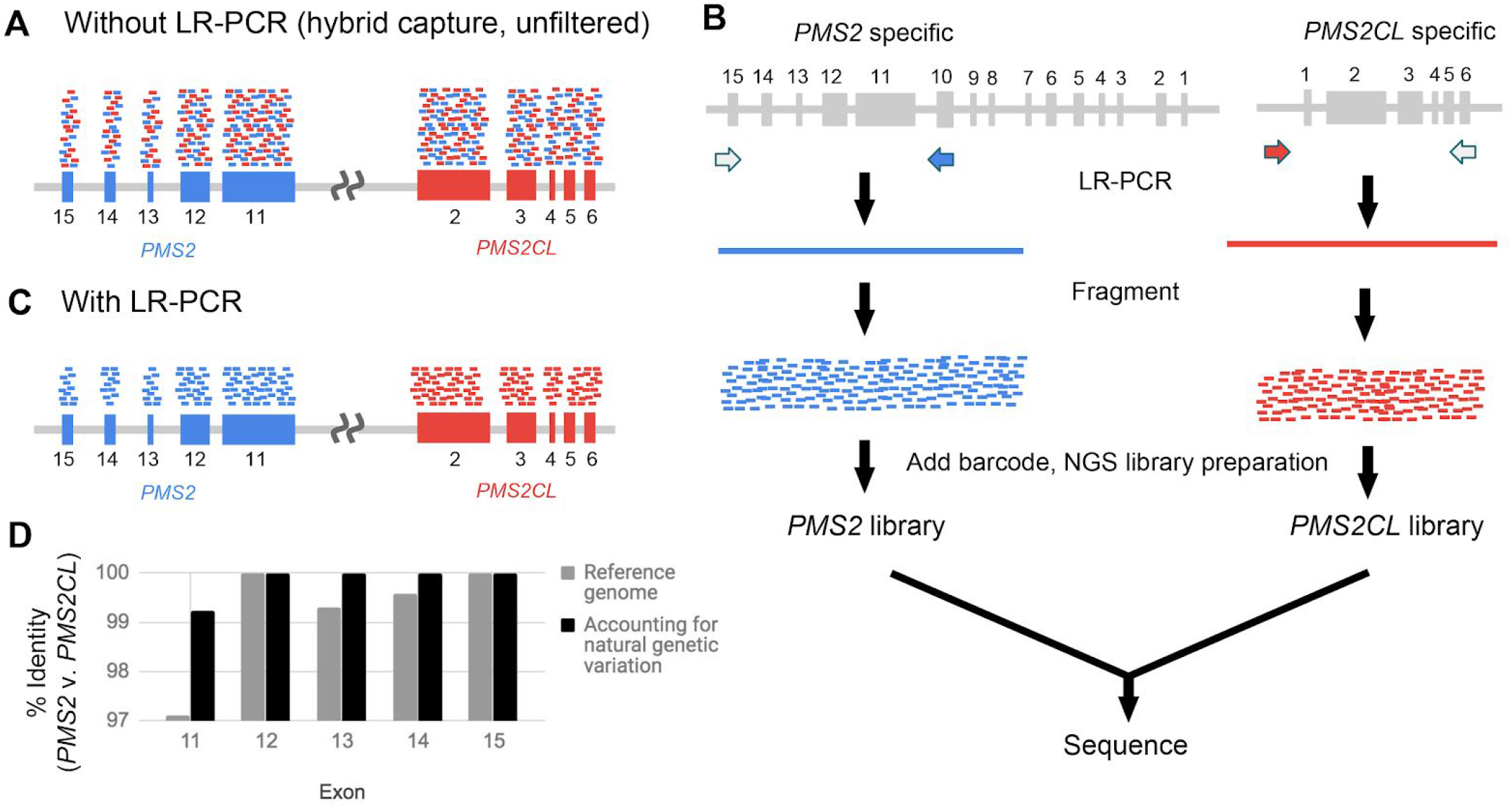
LR-PCR strategy for building a dataset of natural genetic variation in *PMS2* and *PMS2CL*. **(A)** Short-reads from NGS hybrid-capture data that originate from the gene (blue) and pseudogene (red) align to both the gene and pseudogene due to high homology. **(B,C)** Using LR-PCR that is specific to the gene or pseudogene followed by fragmentation and barcoding (B), the resulting short NGS reads can be assigned to the gene or pseudogene (C). **(D)** Percent identity between the gene and pseudogene for *PMS2* exons 11-15 based on the hg19 reference genome (gray) and after accounting for natural genetic variation obtained from LR-PCR samples (black).

Multiple testing strategies exist that can achieve high sensitivity and specificity in the last five exons of *PMS2* [18–20,22,25,26]. Performing LR-PCR, MLPA, and hybrid-capture NGS on each screened sample was presented previously on a small cohort [22], but applying this combination to a large patient population would be resource intensive and complicate workflow logistics. Herman et al. recently presented a method for identifying CNVs (but not SNVs or indels) in the terminal exons of *PMS2* or *PMS2CL* [26]. The method identified samples for follow-up LR-PCR testing to definitively localize the CNV to the gene or pseudogene. The authors noted a CNV false positive rate of 6.8%, meaning that a significant portion of CNV-negative samples would unnecessarily undergo follow-up testing.

Here we present a reflex strategy for detection of SNVs, indels, and CNVs in the last five exons of *PMS2*. Our aim was to have the workflow’s initial testing phase (i.e., upstream of reflex) be sensitive enough to maximize detection of *PMS2* variants and sufficiently specific to minimize reflex burden. The proposed workflow applies hybrid-capture NGS to all samples and LR-PCR/MLPA only as a reflex assay. As the validity of LR-PCR in the last five exons of *PMS2* is established [20,21], we sought primarily to evaluate the performance of the hybrid-capture NGS assay via comparison to LR-PCR results from 299 clinical and cell line samples. We found that the workflow has high analytical accuracy while requiring reflex testing for only 8% of samples. Because our development of this workflow required collection of sequencing data and calculation of variant frequencies from a complicated genomic region with important impact on human health, we have made this information publicly available.

## Materials and Methods

This study was reviewed and designated as exempt by Western Institutional Review Board and complied with the Health Insurance Portability and Accountability Act (HIPAA).

### Study Samples

Table S1 indicates which sample sets were used for particular assays and analyses. Cell-line DNA was purchased from Coriell Cell Repositories (Camden, NJ) (Table S2). Patient sample DNA was extracted from de-identified blood or saliva samples. DNA samples with known positives were a gift from Invitae.

### LR-PCR

DNA was extracted and underwent an additional cleanup via incubation with 1x SPRI beads followed by 80% ethanol wash and elution into TE (10 mM Tris-HCl, 1 mM EDTA, pH 8.0). Approximately 300 ng of eluted DNA served as the template in separate gene- and pseudogene-specific LR-PCR reactions with the following final concentrations: 1x LongAmp Taq Reaction Buffer (New England Biolabs, NEB), 0.3 mM dNTPs, 1 μM of a gene- or pseudogene-specific forward primer, 1 μM of common reverse primer LRPCR_Unv_R (all primer sequences in Table S3), 0.25% Formamide, and 5 units LongAmp Hot Start Taq DNA Polymerase (NEB). Reactions including the gene-specific forward primer PMS2_LRPCR_F yielded a ~17kb amplicon spanning *PMS2* exons 11-15 (the forward primer targets exon 10), whereas use of the pseudogene-specific forward primer PMS2CL_F amplified ~18kb from *PMS2CL* (spans region upstream of *PMS2CL* through exon 6). Thermal-cycling involved initial denaturation at 94°C for 5 min followed by 30 cycles of 94°C for 30 s and 65°C for 18.5 min. Final elongation was 18.5 min at 65°C, followed by a 4°C hold. Quality of LR-PCR amplicons was assessed using 0.5% agarose gel electrophoresis and quantification with the broad range Qubit assay kit (Thermo Fisher).

Two different library-prep strategies were used to prepare LR-PCR amplicons for NGS. In the first, applied to patient samples, LR-PCR amplicons were fragmented by adding 2 μL NEBNext dsDNA Fragmentase and NEBNext dsDNA Fragmentase Reaction Buffer v2 (1x final, NEB) to the remaining LR-PCR reaction volume, and then incubated at 37°C for 25 min. Addition of 100 mM EDTA stopped the reaction, which underwent cleanup with 1.5x SPRI beads, followed by 80%ethanol wash and elution in TE. Fragmentation quality was assessed via Bioanalyzer (Agilent) with the High Sensitivity DNA kit. NGS library prep included end repair, A-tailing, and adapter ligation. Samples were PCR amplified with KAPA HiFi HotStart PCR Kit (Kapa Biosystems) for 8-10 cycles with barcoded primers with the following thermal cycling: initial denaturation at 95°C for 5 min followed by cycles of 98°C for 20 s, 60°C for 30 s, and 72°C for 30 s. The last elongation was 5 min at 72°C, followed by 4°C hold. Library quality was verified via Bioanalyzer with a High Sensitivity DNA kit and the concentration was measured with absorbance via a microplate reader (Tecan Infinite M200 PRO).

The second approach to prepare LR-PCR amplicons for NGS—applied to the 155 cell-line samples—entailed fragmenting and inserting adapters into LR-PCR amplicons via tagmentation. Two duplex adapters were created by annealing single-stranded oligonucleotides: one duplex adapter had the Unv_Tn5_oligo (all primer sequences in Table S3) annealed to Oligo A; the other duplex adapter had the Unv_Tn5_oligo annealed to Oligo B. The two separate annealing mixes included 25 μM of each oligonucleotide in the duplex plus 1x annealing buffer (10 mM Tris-HCl, 50 mM NaCl, 1 mM EDTA, pH 8.0). The reaction was denatured at 95°C for 2 min, incubated at 80°C for 60 min, stepped down in temperature by 1°C every minute until reaching 20°C, and then held at 4°C. Adapters were loaded into the Tn5 enzyme during a 30 min incubation at 37°C with 0.15 units of Robust Tn5 Transposase (kit from Creative Biogene), 1.25 μM of each adapter, and 1xTPS buffer. LR-PCR amplicons were subjected to tagmentation with the Tn5-adapter construct. Tagmentation reactions occurred at 56°C for 10 min in 1x LM Buffer, with 0.5 μL of loaded Tn5 and 1-2 ng of DNA from each LR-PCR reaction. After incubating, SDS (0.02% final) was added to each reaction and incubated for 5 min to dissociate Tn5 from the DNA. Tagmention cleanup with 1x SPRI beads preceded molecular barcoding and amplification via PCR to generate NGS libraries. The PCR reaction included 1 unit Kapa HiFi Polymerase (Kapa Biosystems), 1x HiFi Buffer, 375 μM dNTPs, 0.5 μM of each primer, and the cleaned-up tagmented sample. Cycling started with gap-filling at 72°C for 3 min and followed with 10 cycles of denaturation at 98°C for 30 s, annealing at 63°C for 30 s, and extension at 72°C for 3 min. Cleanup of NGS libraries was performed with 1x SPRI beads.

For patient samples, LR-PCR libraries were sequenced on a HiSeq 2500 (Illumina) in rapid run mode (paired reads, 150 cycles each). For cell line samples, LR-PCR libraries were sequenced on a NextSeq 550 (Illumina) to a minimum depth of 500 reads (single read, 150 cycles).

### Hybrid Capture and Sequencing

Targeted NGS was performed as described previously [7,8]. Briefly, DNA from a patient’s blood or saliva sample was isolated, quantified by a dye-based fluorescence assay, and then fragmented to 200-1000 bp by sonication. Fragmented DNA was converted to an NGS library by end repair, A-tailing, and adapter ligation. Samples were then amplified by PCR with barcoded primers, multiplexed, and subjected to hybrid capture-based enrichment with 40-mer oligonucleotides (Integrated DNA Technologies) complementary to regions common between *PMS2* and *PMS2CL*. NGS was performed on a HiSeq 2500 with mean sequencing depth of ~500x for the whole panel (coverage in *PMS2* is ~1000x). All target nucleotides are required to be covered with a minimum depth of 20 reads.

### Read Alignment

For hybrid-capture data, in order to aggregate *PMS2*- and *PMWS2CL*-originating reads at the *PMS2* locus in the reference genome, paired-end NGS reads were first aligned to the hg19 human reference genome using BWA-MEM [27]. The alignment at *PMS2* exon 11 was filtered to only include reads that overlapped with a site of known difference between gene and pseudogene. Reads that aligned to *PMS2* exons 12-15 and reads that aligned to *PMS2CL* exons 3-6 were partitioned into a BAM file using samtools [28]. The BAM file was converted into two unaligned FASTQ files (each member of the read pair parsed to one of the two files) using Picard (Broad Institute). Each single-end FASTQ file was separately realigned to the hg19 genome allowing for ambiguous alignments and reporting of the top several alignments for each read. The resulting single-end alignments were used to generate a paired-end alignment in the following manner: 1) both single-end reads had the same read name; 2) both single-end reads mapped to the region spanning *PMS2* exons 12-15; 3) both single-end reads aligned within 1000 bp of each other, and 4) when multiple putative pairs met the above conditions for a given read name, the pair with the highest alignment score was chosen. Reads that could not form proper pairs as described above were discarded. The resulting paired-end BAM file contained reads originating from both *PMS2* and *PMS2CL* mapped to the *PMS2* sequence.

For RT-PCR data (described below) and LR-PCR data, NGS reads were aligned to the hg19 genome sequence in which the *PMS2CL* sequence was removed, thereby aggregating genic and pseudogenic reads in *PMS2*.

### SNV and Indel Calling

For the *PMS2* region into which reads from *PMS2* and *PMS2CL* were mapped (see above), SNVs and short indels were identified using GATK 4.0 HaplotypeCaller [29] with the sample-ploidy option set to four. For the diploid *PMS2* exon 11 region, SNVs and short indels were identified using GATK 1.6 [30] and FreeBayes [31]. For diploid SNV calling in the LR-PCR data, GATK 1.6 was similarly used.

### CNV Calling

For short-read NGS data of hybrid-captured fragments, CNVs in *PMS2* exon 11 were determined by measuring the relative NGS read depth at target positions using the algorithm described previously [7]. To call CNVs in *PMS2* exons 12-15 from BAM files in which *PMS2-*and *PMWS2CL*-originating reads were positioned in the *PMS2* sequence (see “Read Alignment” above), two modifications to the CNV calling algorithm were made: 1) the expected wild type copy number was changed from two to four copies, and 2) *p_CNV_* was set to 0.01.

For CNV calling from LR-PCR data, read depth was counted in equal-sized bins (50 bp) that tile the amplicon. Bin counts for each sample were normalized by the median bin depth of the sample; next, each bin’s values were normalized by the median of the bin. The same bins were used for corresponding regions of *PMS2* and *PMS2CL*. The resulting binned and normalized data were searched for CNVs using the algorithm described previously [7]. CNV no-calls were manually reviewed to resolve status as positive or negative.

### CNV Simulations

Single-copy duplications and deletions were introduced by modifying the number of observed reads in one of the CNV-negative samples in a given batch of samples, as described previously [32]. For *PMS2* exons 12-15, where baseline copy-number was four, single-copy deletions and duplications were introduced by subsampling reads to 75% or scaling read number by 125%, respectively. Simulated CNVs were created for every possible contiguous combination of exons in the last 4 exons in *PMS2*. For each CNV size and position, 2186 samples were simulated and tested via the CNV calling algorithm, and sensitivity was calculated as the percentage of the synthetic CNVs that were correctly detected. CNVs were simulated separately in *PMS2* exon 11, which had a baseline copy number of two, because pseudogenic reads were filtered from the genic sequence.

### Tetraploid Indel Simulations

Indels in a tetraploid background (relevant for exons 12-15 of *PMS2*, where gene- and pseudogene-originating reads were remapped) were simulated to better test indel-calling sensitivity using GATK4. Two diploid alignments, at least one of which was previously determined via the Counsyl Reliant HCS panel to contain an indel, were merged to create a tetraploid alignment. If one of the samples had more reads than the other at the site of the simulated indel, reads were binomially downsampled such that each merged diploid sample had approximately the same number of aligned reads. Indels were then called from these synthetic tetraploid alignments using GATK4.

### Variant Curation

For all variants in the last five exons of *PMS2*, variant interpretation was performed in accordance with American College of Medical Genetics and Genomics criteria [33]. The allele-frequency rules were not used because of potentially inaccurate *PMS2* variant identification in public databases.

### MLPA

MLPA was performed according to manufacturer’s protocol (MRC Holland, probemix P008-C1 *PMS2* protocol issued 12/11/17 and MLPA General Protocol issued on 3/23/18). Generally, genomic DNA was covered with mineral oil to reduce evaporation during hybridization and ligation; next, DNA was denatured for 5 min at 98°C and then held at 25°C. Hybridization reagents and probemix were added to the samples and incubated at 95°C for 1 min followed by 16-20 h at 60°C. Probe pairs that bind target DNA at adjacent positions were ligated for 15 min at 54°C and then amplified via PCR for 35 cycles. Amplified probes were mixed with ROX ladder and formamide and then separated on a capillary electrophoresis instrument. Coffalyser software (MRC Holland) normalized *PMS2* probe intensities to those of the reference probes first within each sample and then among samples. Normalized probe intensities of each sample were compared to the average intensities of the reference samples; Coffalyser emitted CNV calls in the region.

### Reflex Rate Estimate

The reflex rate was estimated using SNV-, indel-, and CNV-specific reflex rates from the LR-PCR and hybrid-capture data and subsequently extrapolating to a large cohort size using Markov Chain Monte Carlo simulations with pymc [34].

### Distinguishing Base Analysis

NGS reads from LR-PCR amplicons from *PMS2* and *PMS2CL* were aligned to *PMS2*,and variants were called with GATK UniversalGenotyper. Sites were considered reliable if variants were homozygous for the reference allele in the *PMS2*-specific amplicon and homozygous for an alternate allele in the *PMS2CL*-specific amplicon (as aligned to *PMS2*) in 100% of samples.

### RNA Testing

#### RNA Extraction and Reverse Transcription

RNA was extracted from 33 samples with the Agencourt RNAdvance Blood kit (Beckman Coulter) from 400 μL of whole blood following the manufacturer’s instructions. RNA was extracted from blood tubes no more than seven days after blood draw was performed. Extraction quality was assessed with the RNA 6000 Nano kit (Agilent). RNA was quantified with Qubit HS RNA Assay kit (Thermo Fisher).

RNA was reverse transcribed using Superscript II Reverse Transcriptase with oligo-dT and random hexamers as primers (kit from Thermo Fisher). Reactions were performed as follows: 0.1-1.0 μg total RNA, 1.25 μM of both random hexamers and oligo-dT primer, 0.8 mM dNTPs, and water up to a final volume of 12 μL. Reactions were heated at 65°C for 5 min and then chilled on ice for 5 min. 1x first-strand buffer and 0.01 M DTT were added to each reaction and incubated at 42°C for 2 min. 10 U/μL Superscript II Reverse Transcriptase was added to each reaction and incubated at 42°C for 50 min, then heat inactivated at 72°C for 15 min. A positive control of pooled mRNA (Stratagene, Catalog #750500-41) was used with each reverse transcription reaction.

Following reverse transcription, RNA was hydrolyzed with 2 μL 1N NaOH and heated at 95°C for 5 min. 4 μL of 1 M Tris-HCL pH 7.5 was used to neutralize the reaction for downstream processing. Qubit ssDNA Assay kit (Thermo Fisher) was used to quantify cDNA.

#### PCR

For each sample, two reactions were set up: 1) forward primer PMS2_RNA_F and reverse primer RNA_Unv_R amplified 1.5 kb of *PMS2* from cDNA and 2) forward primer PMS2CL_F and reverse primer RNA_Unv_R amplified 1.5 kb of *PMS2CL* from cDNA (primer sequences in Table S3). PCR reactions contained 1x LongAmp Taq Reaction Buffer (NEB), 0.3 mM dNTPs, 1 μM of each forward and reverse primer, 20-70 ng cDNA, 0.1 U/μL LongAmp Taq DNA polymerase (NEB), and water up to 25 μL. Thermocycling was as follows: 94°C for 5 min, 30 cycles of 94°C for 30 s, annealing at 52°C for *PMS2* and 55°C for *PMS2CL*, 65°C for 2 min, followed by a final extension at 65°C for 10 min and then a 4°C hold. PCR products were cleaned with 1,2x SPRI beads. Amplicons were visualized with a 2% agarose gel or with the DNA 7500 kit (Agilent).

#### Sequencing

50-100 ng of each amplicon were fragmented in 50 μL volumes with a Bioruptor (Diagenode) for 12 cycles, 30 s on and 90 s off. Fragmentation was visualized with High Sensitivity DNA kit (Agilent). All fragmented material was used as input for library preparation. KAPA Hyper Prep kit (Kapa Biosystems) was used for library preparation, and manufacturer instructions were followed. Adapters were diluted to 15 μM for *PMS2* and 3 μM for *PMS2CL*. Nine cycles of enrichment PCR were performed. Samples were quantified using absorbance measurements (Tecan M200), normalized to 10 nM, and consolidated into one reaction. The final library was quantified with qPCR using KAPA Library Quantification Kit (Kapa Biosystems) and sequenced on the NextSeq 550 System (Illumina) for 75 cycles single read with dual indexing.

#### Alignment

Basecall files were converted to FASTQ files using bcl2fastq (Illumina). FASTQ files were aligned using STAR [35].

### Analytical Metrics

Metrics were defined as follows: Sensitivity = TP/(TP + FN); Specificity= TN/(TN + FP). The CIs were calculated by the method of Clopper and Pearson [36]. For SNVs and indels, true negatives were defined as concordant negative results observed at sites found to be polymorphic in our cohort (positions at which we observed non-reference bases in at least one sample).

## Results

### Zero nucleotides can reliably distinguish exons 12-15 of *PMS2* from *PMS2CL*

NGS of short DNA fragments would only be able to identify *PMS2-*specific variants in the last five exons if the fragments themselves could be unambiguously aligned to the gene or pseudogene. To overcome pseudogene interference, unique mapping would rely on the bases that differ between *PMS2* and *PMS2CL*. In the hg19 reference genome, these distinguishing bases are scarce (Fig. 1D, light bars): sequence identity in each of the last five exons of *PMS2* (padded with 20nt of intronic sequence) exceeds 97%, and the differences comprise only 26, 0, 1, 1, and 0 bases in exons 11 through 15, respectively. Further, previous reports noted that natural variation may suppress the reliability of these distinguishing bases represented in the reference genome [17,18].

To test the reliability of the reference genome, we assembled a catalog of natural variation in *PMS2* exons 11-15 and the corresponding regions in *PMS2CL*. We performed NGS on gene- and pseudogene-specific LR-PCR amplicons on 707 of the patient samples in our cohort (Table 1) with diverse self-reported ethnicities (Table S4). We found that 7 of the 26 expected positions in *PMS2* exon 11 had distinct alleles in the gene and pseudogene, making them reliable distinguishing bases. In contrast, for 19 positions in exon 11 and two positions in exons 12-15, the ostensibly *PMS2*-specific alleles from hg19 were observed at least once in the *PMS2CL* LR-PCR data, and vice versa (see Table S4 for allele frequencies). Therefore, after accounting for the natural variation in gene and pseudogene, there are zero reliable distinguishing bases (i.e., 100% sequence identity) in *PMS2* exons 12-15, and seven distinguishing bases in exon 11 (Fig. 1D, dark bars). Together, these data suggest that variant identification via short-read NGS alone could be sufficient for exon 11, but a different approach is required for exons 12-15.

**Table 1:** Summary of samples

| Assay | Samples |
| --- | --- |
| LR-PCR + NGS | 718 patient samples<br>155 cell-line samples |
| Hybrid capture + NGS | 144 patient samples<br>155 cell-line samples<br>3 known positives |
| MLPA | 4 patient samples<br>4 cell-line samples |
| RT-PCR + NGS | 33 samples |

### Reflex workflow to disambiguate variants discovered with short-read NGS

We evaluated the plausibility of a workflow for the 3’ exons of *PMS2* that uses short-read NGS as its foundation and performs reflex testing with orthogonal assays to disambiguate the genic or pseudogenic origin of variants only when clinically needed (Fig. 2A). In the short-read NGS stage of testing, the molecular approach is consistent across the last five exons of *PMS2*:DNA fragments are captured in a manner that is agnostic to their genic or pseudogenic origin by designing capture probes that specifically avoid positions shown to vary between *PMS2* and *PMS2CL* in our LR-PCR data from patient samples (Fig. 2B, purple box).

The workflow employs different bioinformatics strategies for *PMS2* exon 11 and for the group of exons 12-15 (Fig. 2B, blue box). For exon 11, we identified *PMS2*-specific variants by tailoring the read-alignment software to partition reads to *PMS2* or *PMS2CL* based on the gene- and pseudogene-distinguishing bases. By contrast, for *PMS2* exons 12-15, reads are aligned with permissive settings such that each read will align to both its best genic location and its best pseudogenic location (see Methods). For the typical sample with two copies each of *PMS2* and *PMS2CL*, this approach effectively provides read depth in each location corresponding to four copies. To identify SNVs, indels, and CNVs, we adjust the variant calling software such that it anticipates a baseline ploidy of two in exon 11 and four in exons 12-15 (Fig. 2B, blue and green boxes).

**Fig. 2.**
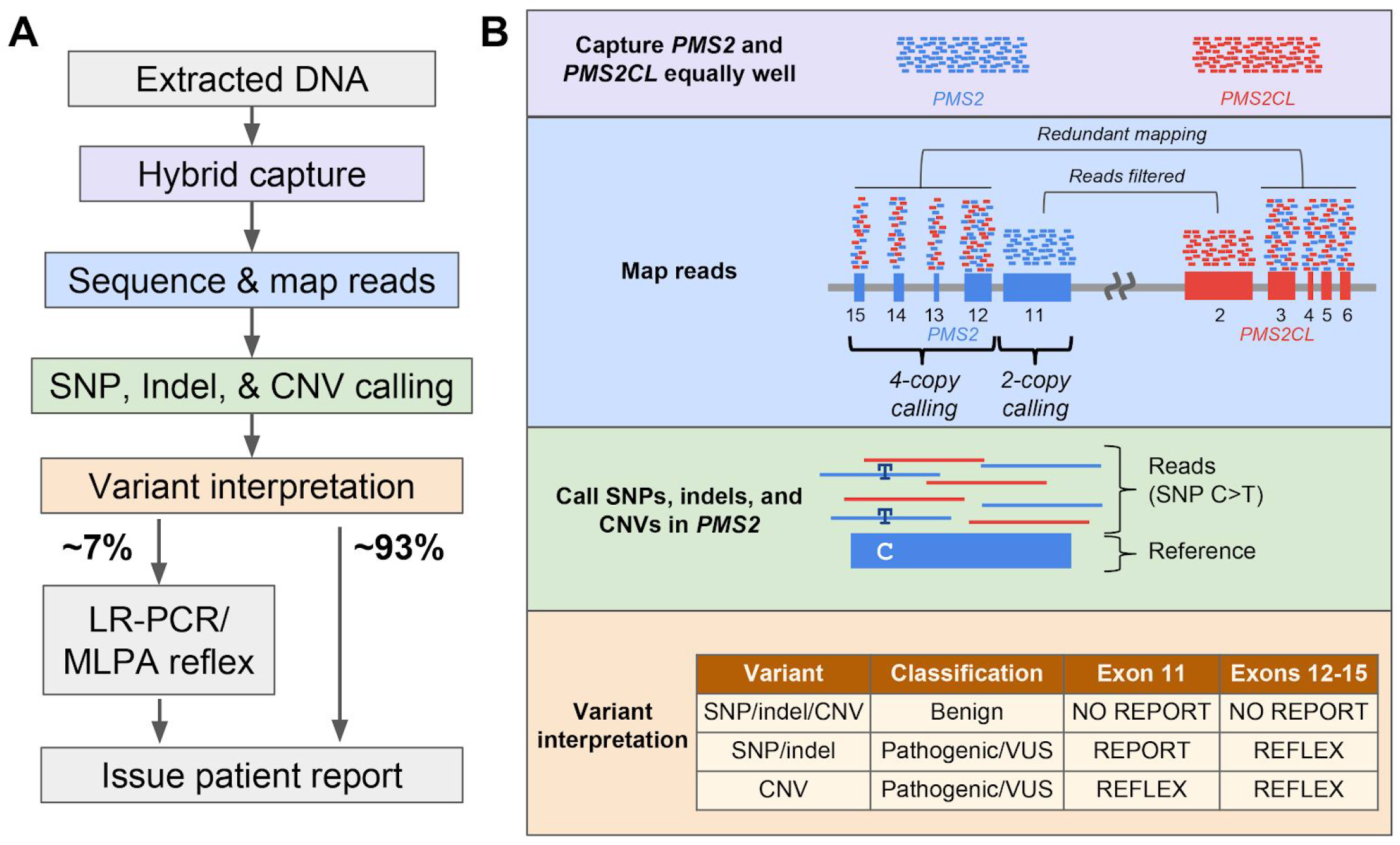
Reflex workflow for variant identification in the last exons of *PMS2*. **(A)** Overview of sequencing and analysis workflow for the last five exons of *PMS2*. Colored nodes correspond to boxes in (B). **(B)** Details corresponding to workflow steps in (A); the details of each box are described in Methods and Results. “No report” means the variant does not appear on patient reports. “Reflex” means the sample is sent for LR-PCR-based disambiguation to determine if the variant is localized to the gene or pseudogene.

Disambiguation via reflex testing is only required for a subset of variants based on their type and clinical interpretation (Fig. 2B, orange box). As such, variant interpretation is performed prior to reflex testing. Benign variants are not reflex tested or reported to patients. Samples with CNVs in any of the last five exons of *PMS2* that are classified as pathogenic, likely pathogenic, or variants of uncertain significance (VUS) undergo reflex testing for disambiguation. Samples with non-benign SNVs or indels in exons 12-15 are reflex tested for disambiguation, but samples with such variants in exon 11 are simply reported without reflex due to unique read mapping in that exon.

Executing the proposed workflow resolves cancer risk associated with the last five exons of *PMS2* for the majority of samples with short-read NGS alone. For each of the 707 patient samples that underwent LR-PCR (Table 1), we performed variant classification on the results and found that nearly 93% could forgo reflex testing. The remaining ~7% would have required subsequent testing to yield confident *PMS2* screening results (Fig. 2A). The SNV- and indel-specific component of this reflex rate was 41/707 (5.8%), and the reflex rates due to CNV calls and no-calls were 2/707 (0.3%) and 1/144 (0.7%), respectively. Using simulations (see Methods), we estimated the reflex rate on a larger cohort of 13,000 patients to be 7.7% (95% CI: 5.4-10.7%). We expect the 0.7% contribution to the reflex rate from samples with CNV no-calls to be an upper-bound estimate because our standard practice of retesting such samples at least once on short-read NGS typically yields a confident negative call (data not shown), thereby avoiding reflex testing. Therefore, we anticipate the overall reflex rate of the proposed workflow would be less than 8%.

### Short-read NGS accurately identified samples needing reflex testing for SNVs and indels

Our proposed reflex workflow is only clinically viable if the short-read NGS test (Fig. 2B) has high analytical sensitivity and specificity for (1) identifying variants in *PMS2* exon 11 and (2) flagging samples that need reflex testing for variants in exons 12-15 with ambiguous *PMS2/PMS2CL* origin. To evaluate accuracy of the short-read NGS testing for SNVs and indels, we compared its results to those observed with LR-PCR for 144 patient samples and 155 cell lines (Fig. 3). Measuring genotype concordance in exons 12-15 required an atypical confusion matrix because short-read NGS genotypes were reported as tetraploid (see Methods), whereas the LR-PCR returned diploid genotype calls for both the gene and pseudogene (Fig. 3A highlights several examples). The matrix includes “Permissible Dosage Errors,” where the presence of alternate alleles is properly detected but the number of alternate alleles is discordant; such errors are deemed permissible because the presence of alternate alleles in short-read NGS would suffice to trigger reflex testing and be corrected. When compared at 1,678 sites with LR-PCR as a truth set, short-read NGS testing had 100% analytical sensitivity and 100% analytical specificity in exon 11 (Fig. 3B), and 99.9% analytical sensitivity and 100% analytical specificity in exons 12-15 (Fig. 3C).

**Fig. 3.**
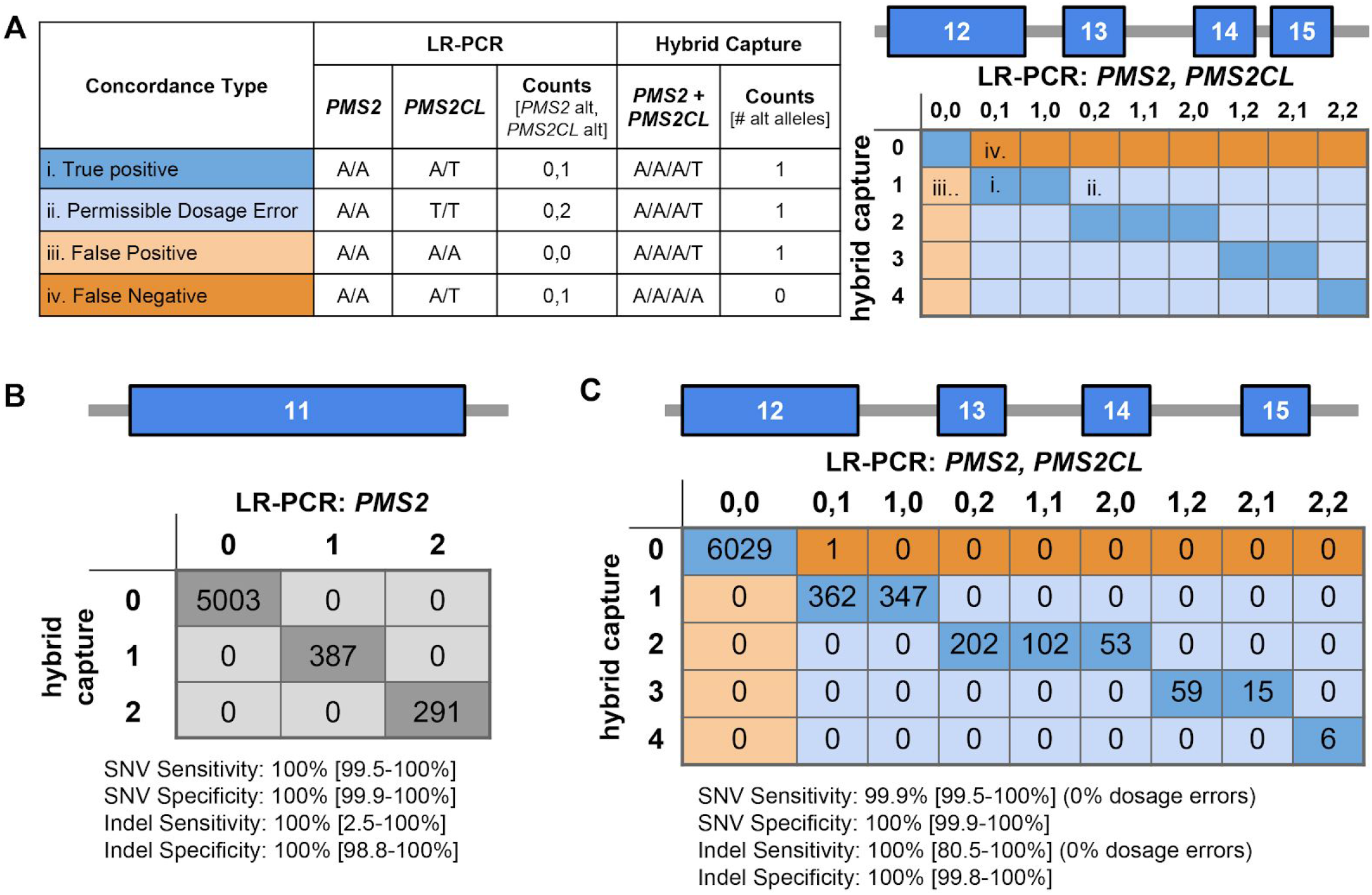
Hybrid-capture and LR-PCR are concordant for SNVs and indels. **(A)** Hypothetical examples to describe the concordance table for comparison of hybrid capture and LR-PCR data. All examples assume the reference base is A and the alternate (“alt”) base is T. (i) Example of a true positive (dark blue) where an alt allele is present in *PMS2CL*. (ii) Example of a permissible dosage error (light blue), where *PMS2CL* is homozygous for the alt allele but hybrid capture only calls one alt allele instead of two. (iii) Example of a false positive (light orange), where only hybrid capture detected an alt allele. (iv) Example of a false negative (dark orange), where an alt allele in *PMS2CL* was missed by hybrid capture. Shaded matrix on the right indicates cells that represent true positives, permissible dosage errors, false positives, and false negatives. Numbers on axes denote the total number of alt alleles in either the hybrid capture data or the *PMS2/PMS2CL* LR-PCR data. **(B)** Diploid SNV and indel concordance for exon 11 of *PMS2*. Numbers on axes denote the number of alt alleles where 0 is equivalent to 0/0, 1 is equivalent to 0/1, and 2 is equivalent to 1/1. 95% confidence intervals in brackets. **(C)** Four-copy SNV and indel concordance for exons 12-15 of *PMS2/PMS2CL*, as explained in (A).

The scarcity of indel calls in our patient cohort and cell lines (17 overall)—coupled with the uncommon usage of variant-calling software in a tetraploid-background mode for a clinical genomics application—motivated a deeper examination of indel-calling efficacy in *PMS2* exons 12-15. We simulated the expected NGS data for samples with a tetraploid genome background populated with indels of different allele dosages (1, 2, 3, or 4 copies). To construct such samples, we merged the diploid NGS data from two samples (at least one containing an indel) in a region of our HCS test other than *PMS2* (Fig. 4A, see Methods). The respective genotypes of the two samples provided an expected genotype of the merged sample: for instance, combining a heterozygous sample (one indel allele) with a homozygous-alternate sample (two indel alleles) would give an expected indel dosage of three. Figure 4B illustrates 99.6% sensitivity for indels in the simulated tetraploid background, suggesting that sensitivity is comparably high in exons 12-15 in *PMS2* where our read-alignment and variant-calling strategy yields a tetraploid background. Because the empirical data in Figure 3C demonstrate 100% specificity for indels in exons 12-15, we did not further evaluate specificity with our simulations.

**Fig. 4.**
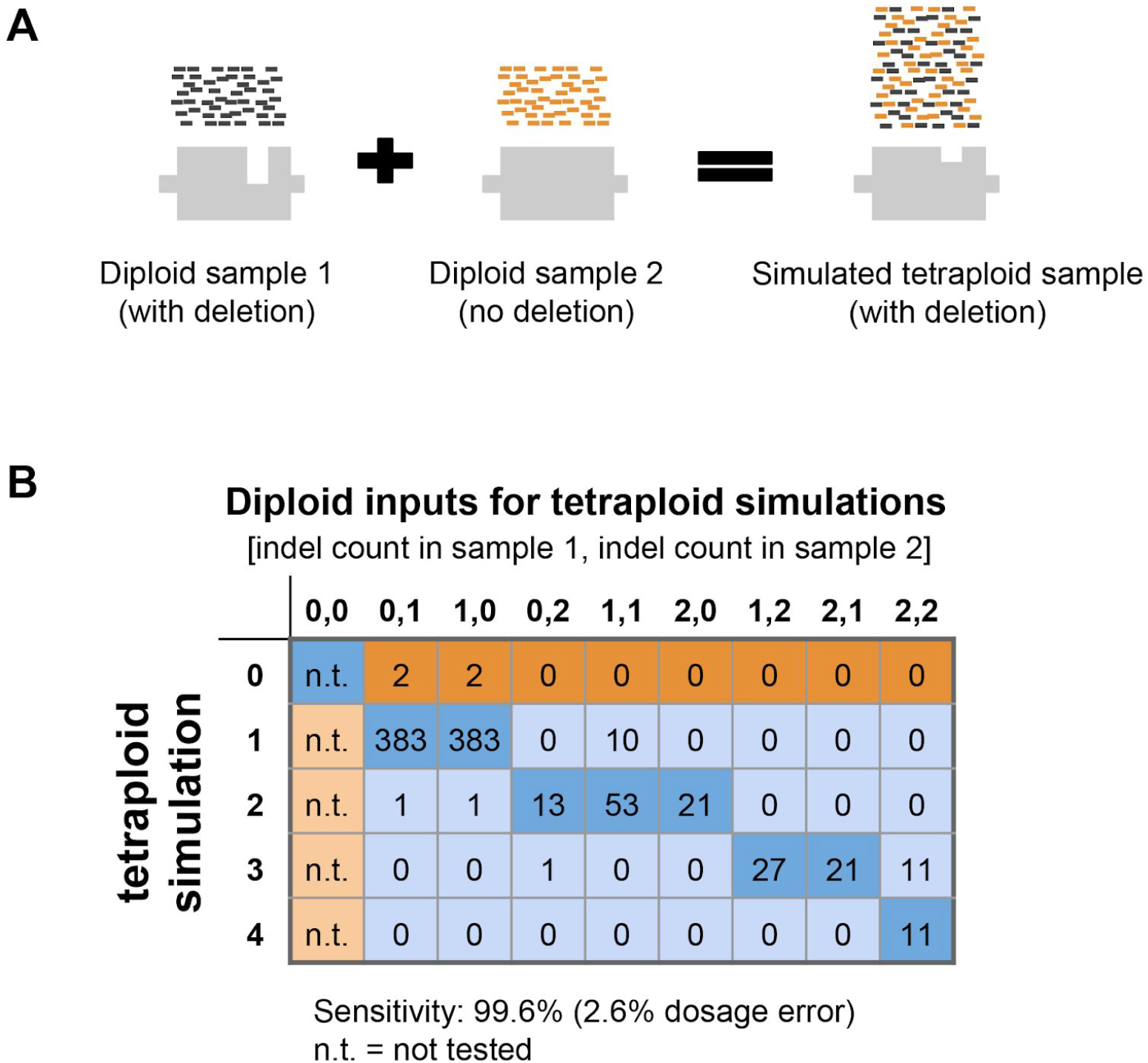
Simulated indels increase confidence in indel sensitivity. (**A)** Schematic of simulating a tetraploid indel by combining sequencing data from two diploid samples. **(B)** Results of tetraploid indel simulations in the same format as Fig. 3A.

In sum, our comparison of SNV and indel calls between LR-PCR and short-read NGS suggests the pre-reflex step of our proposed workflow achieves sufficient analytical sensitivity and specificity to be considered for clinical use.

### Accurate detection with short-read NGS of samples needing CNV reflex testing

To evaluate the sensitivity and specificity of short-read NGS for CNVs in the last five exons of *PMS2*, we tested patient samples, cell lines, known positives, and samples with simulated positives. As with SNVs and indels, we adapted our CNV detection algorithm to use a copy-number baseline of two for *PMS2* exon 11 and four for exons 12-15 (Fig. 2B, blue box; see Methods). The three known-positive samples with CNVs in the last five exons were correctly identified as harboring CNVs encompassing the expected exons (Fig. 5A). We additionally observed a deletion of exon 13-14 in four of the cell lines and one of our clinical samples; for the clinical sample, short-read NGS identified a drop in signal from the tetraploid background (Fig. 5B), MLPA confirmed the presence of a similar deletion (Fig. 5C), and NGS on the LR-PCR amplicons revealed that the deletion was in *PMS2CL* rather than *PMS2* (Fig. 5D). Interestingly, though only one of two copies of this region is deleted in *PMS2CL*, the LR-PCR profile shows a 75% signal drop in the deleted region. We speculate that this arises from preferential amplification of the shorter deletion-harboring allele during LR-PCR. Therefore, although the LR-PCR data were unique in providing disambiguation, the short-read NGS and MLPA data had more readily interpretable copy-number values.

**Fig. 5.**
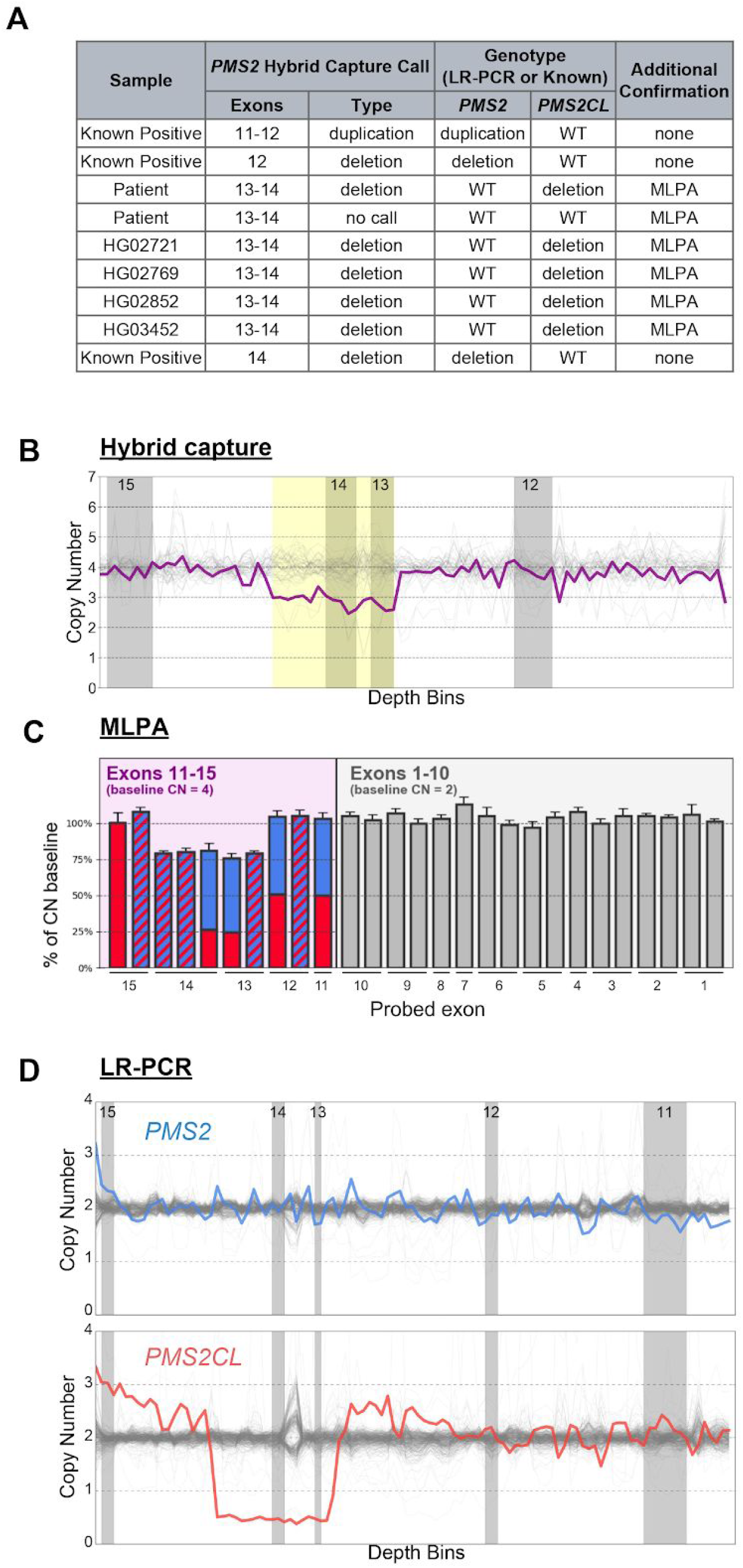
Hybrid capture, LR-PCR, and MLPA are concordant for CNVs. **(A)** All CNVs called in the hybrid capture data and corresponding orthogonal confirmation data. **(B)** Hybrid capture data for the patient sample with an exon 13-14 deletion depicts copy-number estimates across the locus (bins). Gray regions denote the last four exons of *PMS2*. White regions denote introns. Yellow box indicates region of the CNV call. **(C)** MLPA data for the exon 13-14 deletion patient sample. *PMS2*-specific (solid blue), *PMS2CL*-specific (solid red), and *PMS2/PMS2CL* degenerate MLPA probes (blue and red stripes) show the deletion in exons 13-14 of *PMS2CL*. **(D)** LR-PCR data for the exon 13-14 deletion sample depicting copy number estimates across the locus (bins) for *PMS2* (blue, top) and *PMS2CL* (red, bottom). Gray regions depict exons 11-15 of *PMS2* and white regions depict introns as in (B).

Due to the absence of a large catalog of CNV-positive samples, thorough and direct characterization of *PMS2* CNV calling sensitivity with short-read NGS would require blind testing of thousands of samples. Instead, we used sequencing data from the abundance of CNV-negative patients as substrate in simulations that introduce CNVs of given length and location (see Methods). By running our CNV detection algorithm on the 2186 simulated samples, we measured the analytical sensitivity for CNVs ranging from one to five exons in length (Table 2; simulation data on cell-line samples in Table S6). Sensitivity for multi-exon deletions generally exceeded 99.2% and for single-exon deletions was ~89%. Weighing the simulated sensitivities by the observed frequency distribution of CNV length in the last five exons of *PMS2* [21,23,24], we estimate that aggregate CNV sensitivity in this complicated genomic region is 96.7%.

**Table 2:** CNV simulations demonstrate high analytical sensitivity

| Size<br>(exons) | Deletions |  |  |  |  | Duplications |  |  |  |  |  | Overall<br>Sensitivity<br>(weighted) |
| --- | --- | --- | --- | --- | --- | --- | --- | --- | --- | --- | --- | --- |
|  | 1 | 2 | 3 | 4 | 5 | 1 | 2 | 3 | 4 | 5 | Exon<br>11-12 |  |
| **Sensitivity** | 88.9% | 99.2% | 100% | 100% | 100% | 70.0% | 93.8% | 99.3% | 100% | 100% | 100% | **96.7%** |
| **Weights** | 26% | 21% | 8% | 15% | 26% | 0% | 4% | 0% | 0% | 0% | n/a |  |

High sensitivity for CNVs must not come at the expense of low specificity, prompting us to measure the CNV false-positive rate in our large cohort. In our 302 hybrid capture cohort of 302 samples, there was one no-call, which we treat as a false positive. Therefore, sample-level specificity is 99.7% (95% CI: 98.2-100%).

Based on these analyses, we conclude that short-read NGS—as optimized in our described workflow—can achieve >96% sensitivity and >99% specificity for detecting samples with CNVs in the terminal five exons of *PMS2*.

### Gene- and pseudogene-specific variant information for common cell lines

Reference cell lines with known genotypes facilitate development and validation of novel molecular diagnostic methods, yet samples with high-quality genotypes in the *PMS2* region are generally unavailable due to the region’s complicated nature. In the course of developing and testing the workflow characterized above, we performed NGS of both hybrid-capture fragments and LR-PCR amplicons on cell lines where high-quality genome sequences were assembled from whole-genome sequencing with ~30x depth (Illumina Polaris 1 Diversity Panel) or from the Genome in a Bottle (GIAB) Consortium [37,38]. Importantly, Figure S1 shows that the gene-specific genotypes we observed differed from the Polaris and GIAB data (including phased data on GIAB samples; Fig. S1C). In principle, such differences could arise due in part to errors in our data. The concordance between orthogonal hybrid-capture and LR-PCR assays suggests that the genotypes we report here are correct, but as a third orthogonal method, we also genotyped *PMS2* and *PMS2CL* from RNA extracted from 33 of the LR-PCR samples (see Methods). The RNA-derived genotypes were concordant with the LR-PCR data (Fig. S2), strongly suggesting that we elucidated correct gene- and pseudogene-specific genotypes. To aid scientific research and clinical development of *PMS2* and its role in Lynch syndrome, we share the gene- and pseudogene-specific variant information. For patient samples, to share valuable data while being mindful of patient consent and PHI compliance, we provide variant frequencies (Table S4). For cell lines, we share variant frequencies, as well as BAM and VCF files for the LR-PCR amplicons spanning the last five exons of *PMS2* and *PMS2CL* (Table S5 and in ENA accession #PRJEB27948).

## Discussion

Here we show that a reflex workflow starting with short-read NGS and reflexing to LR-PCR and/or MLPA can detect sequence variants in the last five exons of *PMS2* with high analytical sensitivity (>99% for SNVs/indels; >96% for CNVs) and specificity (>99% for SNVs/indels/CNVs). In isolation, short-read sequencing would be incapable of attributing variants to *PMS2* or *PMS2CL*, but it is proficient both at resolving SNVs and indels in *PMS2* exon 11 and at flagging samples with other variants whose origin requires disambiguation via reflex testing. In addition to presenting and testing a comprehensive and plausible workflow, we resolved and have shared the *PMS2*- and *PMS2CL*-specific genotypes and allele frequencies of many hundreds of clinical and cell line samples. Together, the contributions described herein may advance understanding of *PMS2* and facilitate routine screening for Lynch syndrome in HCS offerings.

A high reflex rate after short-read NGS testing (e.g., >10%), while acceptable for the accuracy of a patient’s report, may exert unmanageable logistical overhead on the testing laboratory. The reflex rate has two components—one biological and one technical—each with different sources and constraints. The biological component serves as the floor of the reflex rate: if the assay had perfect analytical specificity (i.e., zero false positives) and clinical accuracy (i.e., correct classifications with no VUSs), then there would nevertheless be a nonzero reflex rate due to the presence of pathogenic variants in *PMS2* exons 12-15 and the corresponding *PMS2CL* regions that need disambiguation. This biological component would, therefore, reflect primarily the integrated population frequency of pathogenic variants across the ambiguous region. The technical component of the reflex rate, by contrast, arises from imperfect analytical specificity and incomplete knowledge of variant pathogenicity. Though higher in our study (99.7%), analytical specificity for CNVs was 93.7% in Herman et al. [26], meaning that the technical component of the reflex rate in that study was at least 6.3% (highlighting the variable nature of the technical component). Also, technical reflex due to VUSs in our workflow was required in 4% of samples, a share that is expected to drop with further screening of *PMS2* and the resulting ability to reclassify VUSs.

There are several laboratory strategies that can yield accurate results for the last five exons of *PMS2*, though each requires quality-control monitoring. These approaches include LR-PCR with Sanger sequencing, LR-PCR with NGS, MLPA (often requires LR-PCR with sequencing to disambiguate if CNV in gene or pseudogene), and a reflex workflow built upon short-read NGS as presented here. A risk of performing LR-PCR alone on all samples is allelic dropout, where one of the alleles amplifies poorly or not at all (e.g., due to a SNV under the LR-PCR primer). Appropriate quality control to mitigate this risk could include examining allele balance at sites across the amplicon: allelic dropout is likely if no heterozygous sites are observed, or if all such sites have allele balance significantly less than 50%. We observed one such sample in our cohort with allele balance of ~7% at SNVs across the amplicon (note that NGS but likely not Sanger sequencing would be able to identify these low-allele-balance SNVs that may be below the Sanger sequencing detection limit); inspection of the hybrid-capture data revealed a SNV under the *PMS2* exon-10 LR-PCR primer (PMS2_LRPCR_F in Table S3), which we broadly observed in 0.13% of patients (i.e., 1 in 769). A shortcoming of monitoring allele balance to flag allelic dropout is that a sample that simply lacks SNVs in the amplified portion of the genome could undergo extensive follow-up characterization without actually being spurious. Ultimately, an asset of a workflow incorporating multiple orthogonal methods—e.g., both LR-PCR and hybrid-capture-based NGS (in parallel or in a reflex arrangement)—is that the different datasets facilitate reconciliation of complicated genotypes.

We attempted to mitigate several potential limitations of our proposed workflow. For instance, our short-read NGS approach in *PMS2* exons 12-15 operates variant-calling software with the assumption of a tetraploid background, obviously unusual in a human clinical genomics setting (GATK supports tetraploid calling, but reports of its efficacy in this mode are scarce). Importantly, there was high concordance between short-read-generated SNV calls in the four-copy background and the combined genotypes detected using gene- and pseudogene-specific LR-PCR. A dearth of cell-line or patient samples with indels or CNVs in exons 12-15 also complicated the ability to assess performance of detecting these important variants. In silico simulations enabled generation of 940 indel-positive and 2186 CNV-positive samples in a tetraploid background, and variant calling on these simulated samples revealed high sensitivity. Finally, despite examination of the workflow’s variant-calling accuracy on hundreds of samples, the assay would still require validation before being demonstrated suitable for clinical use.

Although we measured the sensitivity and specificity of the proposed workflow, its potential impact on cost and turn around time (TAT) of the test were not explored here. The impact on cost and TAT depends greatly on how a laboratory decides to implement the reflex-testing workflow. For instance, TAT could be minimized by running the LR-PCR reactions as soon as samples arrive and then only perform NGS of the amplicons upon flagging by the short-read NGS analysis. But, this approach would incur the cost of generating amplicons in >90% of samples that would not need further testing. By contrast, cost is minimized by only doing LR-PCR with NGS on relevant samples after the short-read NGS step, but this approach could increase TAT for those samples. These considerations are ultimately important because TAT and cost impact the utility and accessibility of HCS. It will be exciting to see if future technical developments enable targeted long-read sequencing, as this advance would facilitate clinical-grade testing of highly homologous regions of the genome.

## Conclusions

Screening for pathogenic variants in the last five exons of *PMS2* is technically challenging. High homology between *PMS2* and *PMS2CL* complicates identification of gene-specific variants with short-read NGS alone, and reference cell lines that typically aid assay development and validation are not accurately genotyped in public databases. To help overcome these limitations, we have characterized a reflex workflow that achieves high accuracy and publicly shared the gene- and pseudogene-specific raw data, genotypes, and variant frequencies in widely available cell lines.

## Abbreviations

HCS: :Hereditary cancer screening;
NGS: :Next-generation sequencing;
LR-PCR: :Long-range PCR;
RT-PCR: :reverse transcription PCR;
SNV: :Single-nucleotide variant;
indels: :Short insertions and deletions;
CNV: :Copy-number variant;
MLPA: :Multiplex ligation-dependent probe amplification;
NEB: :New England Biolabs;
VUS: :Variant of uncertain significance;
GIAB: :Genome in a Bottle;
TN: :True negative;
FN: :False negative;
TP: :True positive;
FP: :False positive;
TAT: :Turn around time

## Declarations

- Ethics approval and consent to participate: The protocol for this study was reviewed and designated as exempt by Western Institutional Review Board.
- Consent for publication: Not applicable
- Availability of data and material: BAMs for the cell-line LR-PCR data can be found in the European Nucleotide Archive (accession #PRJEB27948). Corresponding VCFs of these samples are included as a supplemental file.
- Competing interests: All authors are current or former employees and equity holders of Counsyl.
- Funding: The study was funded by Counsyl.
- Authors’ contributions: GMG, PVG, MRT, DHH, CSC, JRM, GJH, and DM designed the study. GMG, PVG, MRT, LS, IEW, LMM, and RGC collected and analyzed the data. GMG, PVG, MRT, LS, IEW, LMM, GJH, and DM wrote the manuscript. All authors read and approved the final manuscript.
- Acknowledgement: We are grateful to Kaylene Ready, Kristin Price, Leslie Bucheit, Peter Kang, Kevin Haas, Kevin Iori, Eric Evans, Krista Moyer, Becca Mar-Heyming, Kerri Hensley, Megan Judkins, Christine Lo, Eric Olson, Kyle Beauchamp, Kristjan Kaseniit, Jeffrey Tratner, Henry Lai, Carly Paul, Pranav Sharma, Victoria Brewster, Irina Ridley, Harris Naemi, and Gabor Brasnjo for support of this manuscript. We thank Invitae for providing CNV-positive samples.

---

**Table S1: Samples and cell lines used in particular assays and analyses**

**Table S2: Cell-line samples included in study**

**Table S3: Oligos and primers used for LR-PCR, RT-PCR, Tn5 adapters**

**Table S4: Allele frequencies from 707 LR-PCRs**

**Table S5: Allele frequencies from 155 GIAB and Polaris LR-PCRs**

**Table S6: Simulated CNV Sensitivity in Cell Line Samples**

**Fig. S1.**
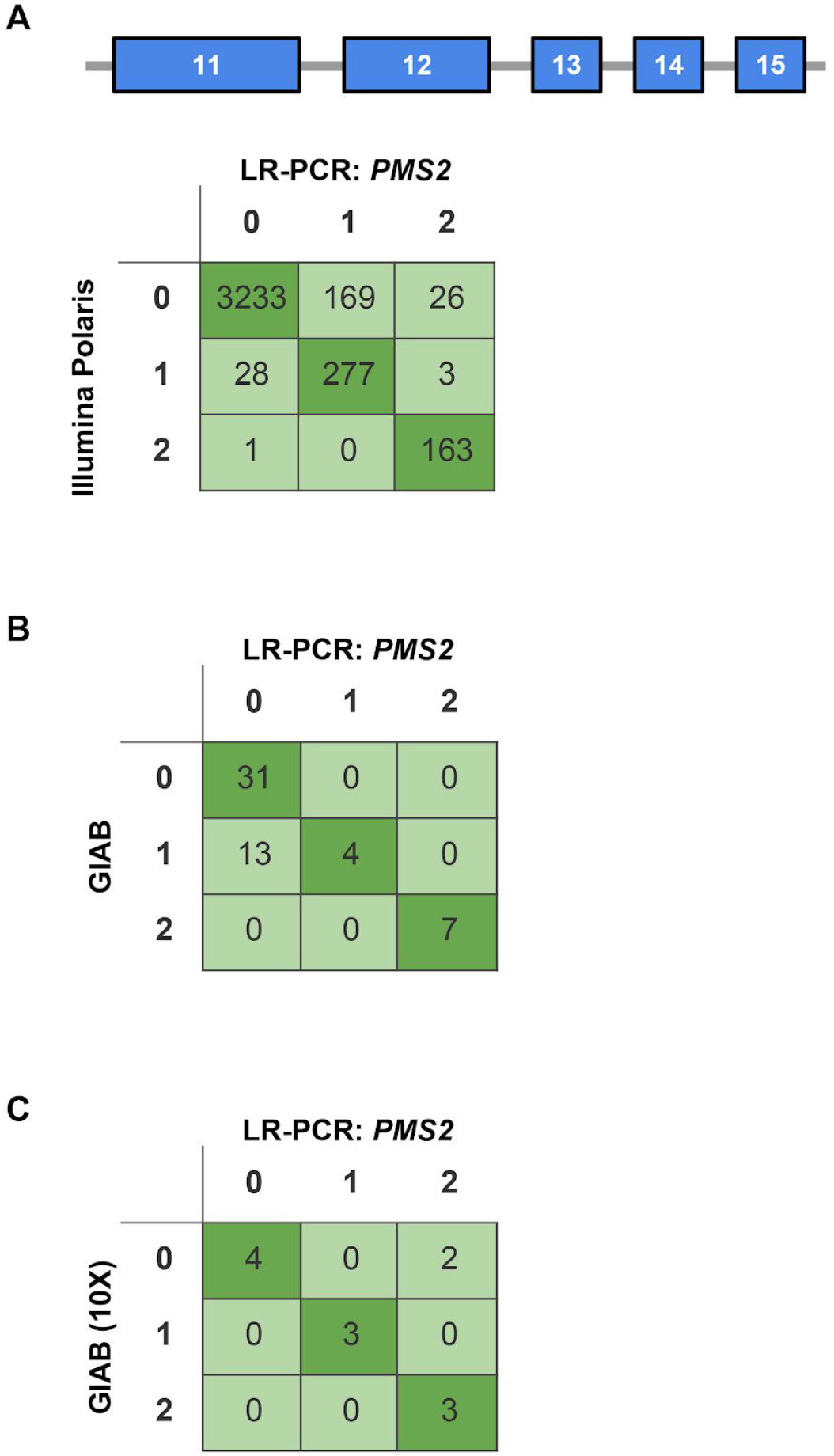
*PMS2* exons 11-15 reference genotypes (from Polaris and GIAB) are inconsistent with *PMS2* LR-PCR. **(A)** Concordance between LR-PCR variant calls and Polaris variant calls. **(B)** Concordance between LR-PCR variant calls and the GIAB multisample call set (including high confidence and filtered variant calls) for all five GIAB samples. **(C)** Concordance between LR-PCR variant calls and the 10X Genomics haplotype call set available for four GIAB samples.

**Fig. S2.**
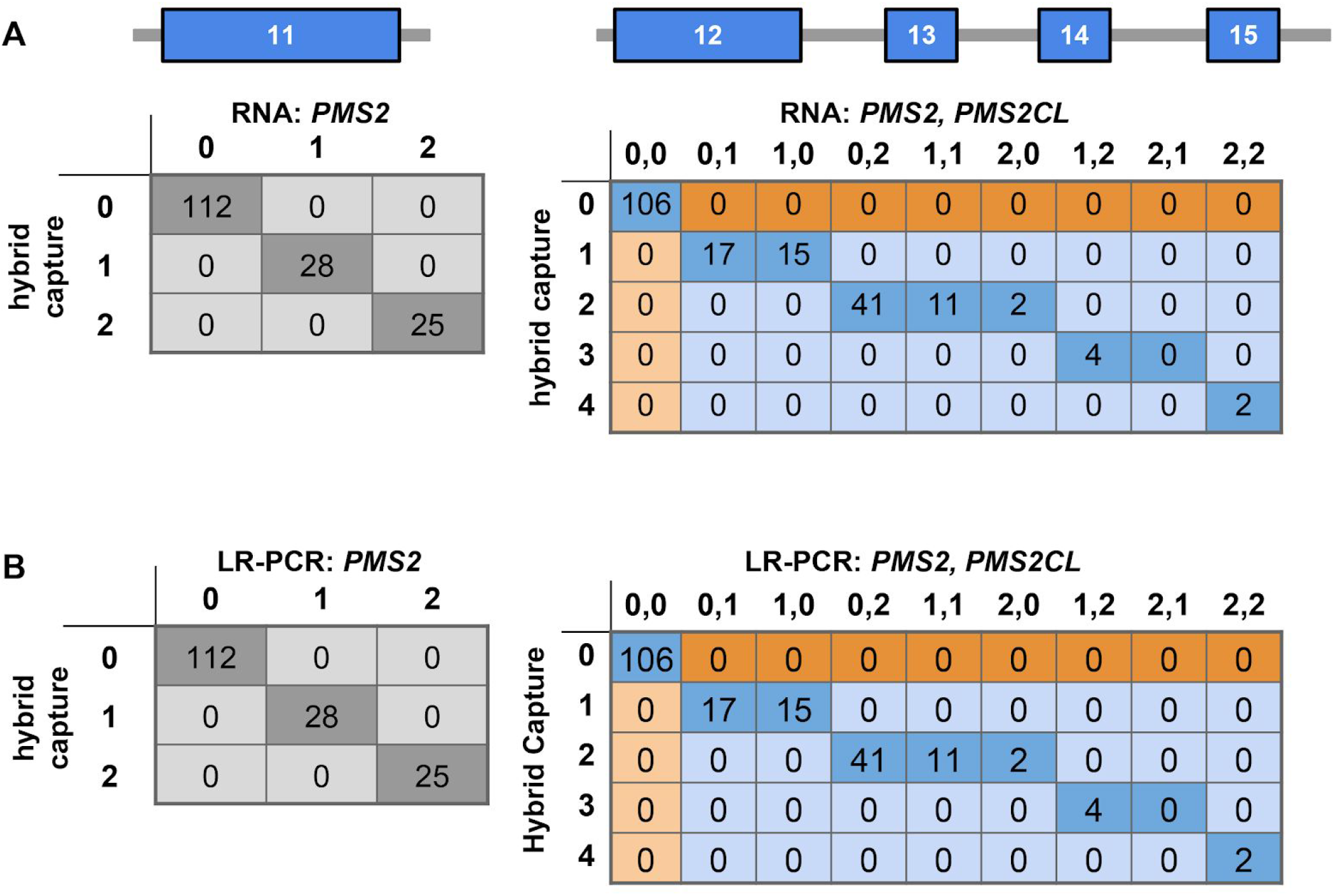
RNA data corroborate hybrid capture and LR-PCR data. **(A)** Concordance between hybrid capture data and RT-PCR for *PMS2* and *PMS2CL*.**(B)** Concordance between hybrid capture data and LR-PCR for *PMS2* and *PMS2CL*.

